# Elastic Multi-Scale Mechanisms: Computation and Biological Evolution

**DOI:** 10.1101/111666

**Authors:** Juan G. Diaz Ochoa

**Affiliations:** PMQD GmbH, Pelargusstr. 2, 70180 Stuttgart, Germany

**Keywords:** Systems Biology, Elastic Mechanisms, Computational Theory, Turing Machines, Evolution, Open Ended Evolution

## Abstract

Explanations based on low level interacting elements are valuable and powerful since they contribute to identify the key mechanisms of biological functions. However, many dynamic systems based on low level interacting elements with unambiguous, finite and complete information of initial states generate future states that cannot be predicted, implying an increase of complexity and open-ended evolution (oee). Such systems are like Turing machines, that overlap with dynamical systems that cannot halt. We argue that organisms find halting conditions by distorting these mechanisms, creating conditions for a constant creativity that drives evolution. We introduce a modulus of elasticity to measure the changes in these mechanisms in response to changes in the computed environment. We test this concept in a population of predators and predated cells with chemotactic mechanisms and demonstrate how the selection of a given mechanism depends on the entire population. We finally explore this concept in different frameworks and postulate that the identification of predictive mechanisms is only successful with small elasticity modulus.

## Introduction

The combination of both data science and network theory has become prominent to model and predict outcomes in many complex systems, from biology and medicine to society. Data produces the necessary knowledge to deduce mechanisms responsible of interactions between different elements, and allow us to conclude if these interactions represent a physical or at least a causal relation between them. These mechanisms are helpful to identify basic principles of biological functions, helping to better approach the Cartesian vision of Biology: "I should like you to consider that these (biological) functions follow from the mere arrangement of the machine’s organs every bit as naturally as the movements of a clock or other automaton follow from the arrangement of its counter-weights and wheels" (Descartes, Treatise on Man, p.108).

Despite this, there is still not a well-established and unified mathematical formalism providing an essential description of complex biological systems. Only network theory has become the best candidate to derive the fundamental basis of this formalism, in part because it allows the mathematical implementation of mechanistic explanations.

The obvious relevance of this methodology is the possibility to write predictive models, allowing the extrapolation from one to other system (translation) as well as the extrapolation in the future (prediction). For instance, molecular biology is essentially reductionist, since it assumes that “explanations that come from lower levels are better than explanations that come from higher levels” (see (Tabery et al., 2005)). Therefore, “biological theories need to be grounded on molecular biology and on physical sciences, for it is only by doing so that they can be improved, corrected, strengthened, and made more accurate and more adequate and complete” (Rosenberg, 1997). Biological theories are then represented by models, which are usually constructed on the base of network theory. Ideally these networks can be experimentally validated, and then be used for extrapolation (in time to predict future events or between different target systems, for instance in drug tests from animal models to humans).

But it is impossible not to feel some discomfort in this development. An extrapolation based on a model requires a perfect and complete mathematical description. This goal can be reached in physical sciences, when perturbation sources are controlled or well characterized, allowing the discovery of fundamental interaction mechanisms. But in complex systems we continuously experience the limits of this methodology, since we are continuously exposed to open and evolving systems.

It is usually granted that the context of an organism and its evolution takes place after long time periods, but in practice it is difficult to know how fast this evolution is. For instance, in medicine: “disease cannot always be predicted with certainty, and health professionals must identify and modify risk factors. The common unidimensional “one-risk factor to one-disease” approach used in medical epidemiology, however, has certain limitations “(Ahn et al., 2006). This concept is the origin of epidemic paradoxes, i.e. spread of apparent diseases in healthy populations simply by adjusting the threshold values of characteristic physiological measurements-like levels of biomarkers- (Ahn et al., 2006). Not only in risk assessment, but also in the identification of evolution this concept is blind. In pharmacology we observe an intermittent adaptation of organisms to substances after long administration times, which are much shorter than time spans for biological evolution (see (Peper, 2009) for a mathematical model considering interactions with the environment)^1^.

These facts challenge the construction of models. In effect, the myriad of possible interactions motivates a continuous update in the information registered in data bases. Thus, while some canonical pathways are well known, many other interactions, and possible variations, are still unknown and must be constantly updated when these mechanisms are reconstructed.

These variations are rooted in many possible ways to respond to the environment. The interconnection between different scales is a central concept that, despite its growing relevance in sciences, is usually ignored (Ellis, 2012), and is related to the autopoietic character of biological systems. A hierarchic construct seems to be a first good approach, if small structures are involved in collective interactions that drive self-organization producing larger structures. But this organizational characteristic is not enough to understand how “assimilation” and “accommodation” works for systems included in larger organisms (Bitbol and Luisi, 2004).

Like condensed matter (Castelvecchi, 2015), and its following concepts from computational theory, micro processes in biological organisms are fundamentally incomplete and undecidable, implying that “there is more than a crude metaphor behind the analogy between cells and computers” (Danchin, 2009).

Only when these processes are included in the organism is completeness achieved, allowing the evaluation of predictions by using a mathematical formalism (which is the background for systems biology). We argue that to get this decidability, biological processes change their fundamental structure, changing also its relationship with the environment. This implies that systems not simply emerge or are merely autopoietic, but model themselves, in the spirit of basic cognitive notions as “assimilation”, “integration”, and “accommodation” to an environment (Bitbol and Luisi, 2004). To this end, we introduce an “elasticity” modulus to measure how strong the deformation of these mechanisms and cross-scale interactions are. We use this module as a measure, and demonstrate its use in a toy model for chemotaxis, complementing current methods based on network theory. Finally, we discuss this theory in the framework of systems theory and explore potential applications.

## Vesicles and completeness

Consider for instance the formation of vesicles, starting from a relatively static aqueous system (vesicle) formed by a surfactant *S.* Here a highly lipophilic precursor of S, indicated as S–*S*, binds to the boundary of the vesicle and is hydrolysed there. The vesicle grows, and eventually divides into two or more thermodynamically more stable smaller vesicles. The more vesicles are formed, the more *S–S* is bound, forming additional vesicles, making the process auto-catalytic. Since the entire process of hydrolysis and growth takes place because of and within the boundary, the vesicle can be seen as a simple self-reproducing, autopoietic system” (Bitbol and Luisi, 2004).

These simple vesicles contain internal reactions mechanisms, as for instance when “the membrane (***S***) is formed from the molecule ***B*** through a process characterized by a generation velocity *v*”. Then, ***S*** decays with velocity *v_deg.* Additionally assume that the precursor metabolite **A** enters from the environment, and that ***B*** decays into ***C***, which is eventually expelled.” This cycle can continue at infinite if the process is homeostatic and has access to energy sources able to maintain it. At this point all these processes are mechanistic, and can be described using simple ordinary differential equations (see figure 1).

**Fig. 1:**
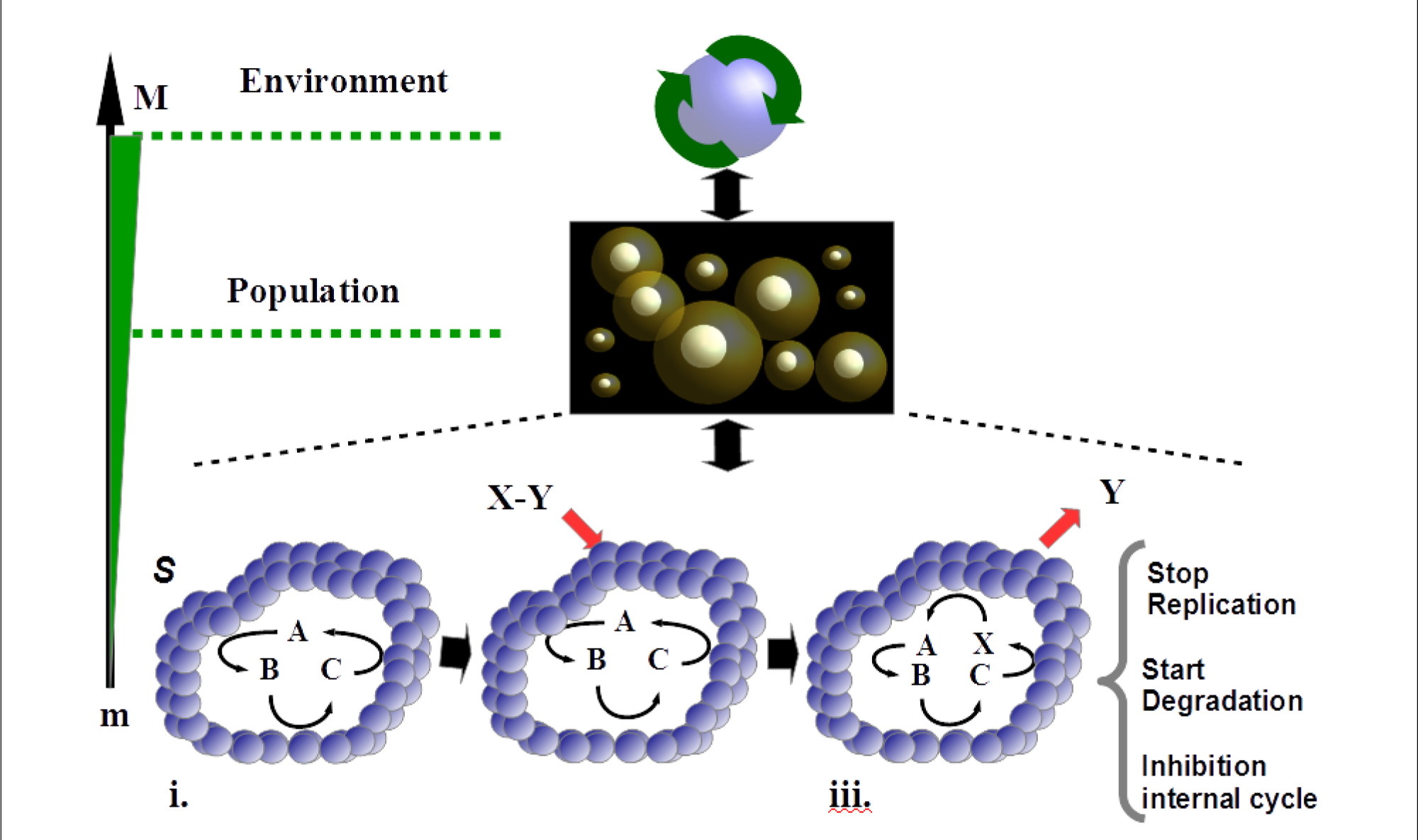
Example of a vesicle with a membrane formed by a cycle generated with the molecules A, B and C (i) that allows the vesicle replication. After interaction with the nearest population as well as with the environment a pair of molecules X-Y is absorbed(ii); thereafter the molecule X is linked into the cycle (iii). This interaction can trigger important “decisions” in the vesicle concerning its maintenance or degradation, or the inhibition of the internal cycle.

Regarding the dynamics of the vesicle, the work of Maturana and Varela is useful to understand how the interaction between an autopoietic unit and the environment can change by considering basic concepts of cognition (Maturana and Varela, 2004). Indeed, there is adaptation whenever “there is: (a) an environmental cause, in the example of the vesicles an outer molecule ***X-Y***; (b) a resulting effect from the unit, here the inception of the molecule ***X*** in the cycle and the release of the metabolite ***Y***; and (c) an adaptive virtue of the effect (Bitbol and Luisi, 2004), the inhibition of the vesicle reproduction, triggering of a degradation process. This cycle can repeat ad infinitum.

The fact that the bounded molecules ***X-Y*** may trigger or inhibit internal processes (from the unit) implies the introduction of an *external decision* about the next state of the vesicle, regarding the population of other vesicles and the environment. The population with the initial set of reactions (square symbol in figure 1) surely fulfills all thermodynamic constraints, and is associated to a high fitness at the population level. But these optimal conditions do not help the vesicle to decide about, for example, when to stop or continue the replication process.

In this example, biological processes cannot make decisions “by themselves” based only on mechanistic (and physical) notions at the micro level. We suspect that this issue is common for many systems in biology and other sciences. For instance, for 2D lattices of atoms “the undecidability ‘at infinity’ means that even if the spectral gap is known for a certain finite-size lattice, it could change abruptly — from gapless to gapped or vice versa — when the size increases, even by just a single extra atom. And because it is “probably impossible” to predict when — or if — it will do so, it will be difficult to draw general conclusions from experiments or simulations “(Castelvecchi, 2015) (Cubitt et al., 2015). This is a remarkable result since it extends concepts from the theoretical computation theory into physics.

We postulate that internal properties in vesicles are like changes in internal states of atoms, such that populations of vesicles behave like a Turing Machine making them unable to halt by themselves. Therefore, in this framework biological processes require assimilation and accommodation.

Thus, our hypothesis is that micro biological systems are inherently incomplete, and that assimilation and accommodation help them to be complete and decidable. For instance, if the population of vesicles has enough energy then they can eventually continue reproducing (like an immortal cancer cell in a tumor), which is nothing different to an infinite iteration of a Turing machine. Only when this unit belongs to an organism this process will be decided (i.e. can continue forever or can be stopped) according to the functionality of the population as an organism.

## Open ended evolution (oee), computability and biological mechanisms

### Mathematical background

The goal of mathematical descriptions of a biological system is to compute its change and evolution, assuming that it is like a deterministic or stochastic system that can be mathematically described with continuous real functions or discrete states that evolve according to certain rules, like for instance Boolean rules.

Alternatively, biological systems are like computational machines that process input information to compute next states. Thus, this category is not a representation, but assumes that de facto biological systems are closer to a computation than classical dynamical system. In the last case, “an object x is computable if it can be described by machines like a Turing machine (Turing, 1936); for example, if there exists a Turing machine that produces x as an output.”(Hernández-Orozco et al., 2016).

Despite recent efforts to demonstrate the link between dynamical systems and computability theory (Baescu et al, 2016), for instance by showing that analytical and real functions are computable (either with classical or quantum computational methods), there is no rigorous proof showing the equivalence between computational theory and dynamical systems theory. We opt here to assume that biological systems are like computational machines that overlap with a dynamical system. The examples shown in the previous section are dynamical systems with unambiguous, finite and complete information of initial states, i.e. computable systems. According to the formalization of Turing and Church, for these computational systems there are future states that cannot be predicted (Turing, 1936).

As has been shown by Hernández-Orozco et. al. (Hernández-Orozco et al., 2016) decidability imposes limits to the growth of complexity, and for this reason oee, particularly non-decidability, are required to understand the complexity growth in biology. However, increase of complexity in cooperating organisms across scales can be life threatening, since processes must halt while a constraint in the complexity growth is retained (for example apoptosis in cells stops the life cycle of the cell, sustaining the correct function of large organisms by maintaining a relatively constant population of cells). Since the behavior of the organisms depends on its adaptation and survival regarding an environment, these biological computable systems also depend on an effective representation of the environment.

The computation *p* of 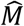, 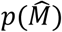 determines how the micro states change in time, i.e. *p:σ_i,t_ →σ_i,t+1_,* where 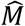 is a structure where information is stored, and 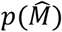 the set of rules to estimate the transition from one input state to the next. Therefore, *p* codifies the rules required to estimate the next state *σ_i,t_* using the information stored in the network structure 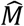. The computation 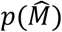 is apparently isomorphic to the dynamical description of the system. However, from the rigorous point of view, both are completely different mathematical objects (Baescu, 2016). When this computation represents the environment *E*, then (see example 1 in table 1)

**Table 1.**
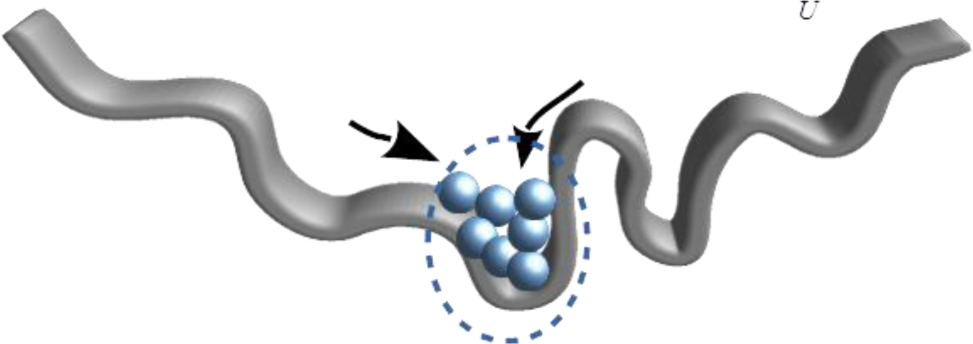
*How environment and control functions are internally represented in elements belonging to a system*

Example 1: Homeostasis is related to the optimization of landscape *U* in a control function (for instance fitness) close related to the environment. Each element (small spheres), as well as the whole organism (sphere population enclosed in a dotted circle) explore with their trajectories *Γ* their environment *E* and are constrained to the optimization ({*max, min*}*U*) of this landscape.

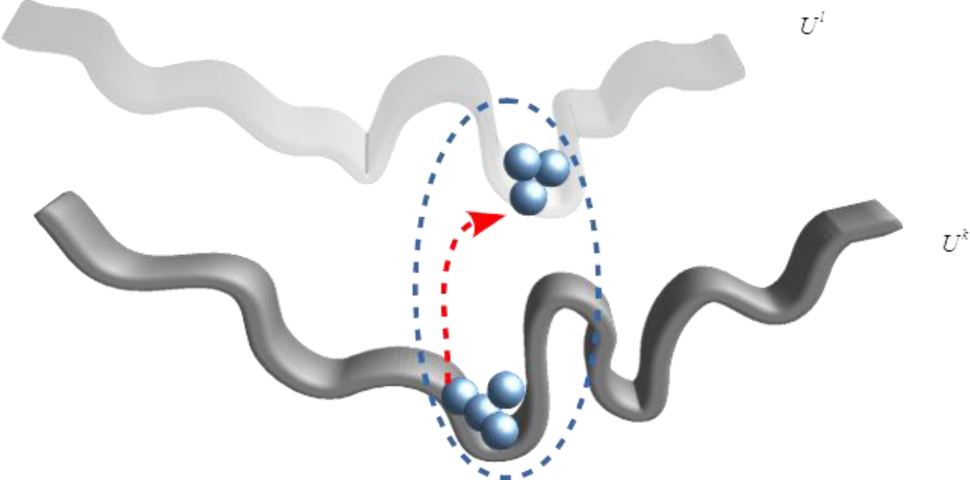

Example 2: The control function *U^k^* is locally optimized, but the entire population is eventually constrained to a different control function *U^l^.* The gap between trajectories in eq. 3 depends on the *u^l^* accommodation and assimilation of the organism (population) to these environments, or equivalently to the use in the organism of compensating mechanisms, such that biological functions are decided.

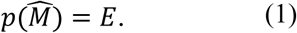

Following the definitions introduced by Hernández-Orozco et al. (Hernández-Orozco et al., 2016), the information of the organism is codified in a structure 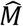, which contains several microstates ordered in such way they can have a response to external inputs. This structure can be represented by a network, such that 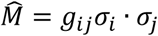 is the connectivity factor of the information of microstates *σ_i_* and *σ_j_.*

“In weakly convergent systems, the program 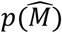 represents an organism, a theory or any other computable system that use the structure 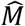to predict the behavior of *E*” (Hernández-Orozco et al., 2016),. If this representation cannot be attained in a certain time period, then the system is *non-adapted,* or the theory is *non-useful.*

Since this theory overlaps computational theory with dynamic systems, the computation of the environment *E* means the computation of the trajectories Γ representing the environment *λ*, i.e. *E^λ^ = Γ^λ^* (for this notation see Wang et al. (Wang et al., 2012b)). This process can be seen as the search of food in a grid, or the expression of proteins in a genetic network expressing a phenotype.

If the environment is dynamic, i.e. *E*(*δ_t_*) for a sequence of times *δ_1_*…*δ_t_*…, then there exists a computation *p_t_*, such that 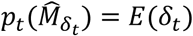 or in other words there exist a trajectory Γ(*δ_t_*). According to the theorem 13 in Hernández-Orozco (Hernández-Orozco et al., 2016), if a series of representations of the environment *p: t* ↦ *p_t_* is computable, the function *δ:t* ↦ *δ_t_* is computable, and the descriptive complexity of the system is bounded. On the other hand, if the sequence of times *δ:t* ↦ *δ_t_* is non-computable, then the system is non-computable, i.e. the environment is non-computable. Thus, there is no way to decide if Γ (*δ_t_*), with this sequence of times, represents the environment, implying its growth of complexity. The definition above (Lemma 11 and theorem 13, corollary 14 (Hernández-Orozco et al., 2016)) represent the formal basis of open ended evolution (oee).

For non-cooperative organisms, *p_t_* cannot find computable conditions to represent its dynamical environment, increasing the complexity of the system. Such organisms are sensitive to *E*(*δ_t_*) but are in general blind to further variations of the environment. However, organisms can cooperate with other organisms and the environment, and recognize that the increase of the complexity can be life threatening (for instance non-apoptotic cells in cancer tissues). For this reason, we argue that systems not only own oee, and that they look for computability conditions.

Computability can be imposed if more than one (dynamical) environments exist and *λ* are computed. Under this condition, the system explores a family of trajectories *Γ^λ^*(*δ_t_*) searching for computability conditions in the environment, and generating in this way a biological function. This is a plausible condition in multiscale systems (cells, organelles, organs, etc.), such that

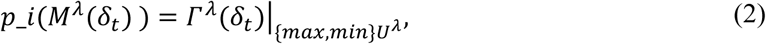

where *Γ^λ^*(*δ_t_*)|_{*max,min*}*U^λ^*_ is the trajectory of the microstates constrained to the optimization (maximization or minimization) of the control function *U^λ^* related to the environment *λ,* which simultaneously belongs to a family of environments *λ = 1…* Λ (see table 1).

This condition implies a distance between two trajectories belonging to two different environments *k* and *l (k* and *l ϵ λ = 1…* Λ), which is the absolute value of the mean distance between the points of two trajectories

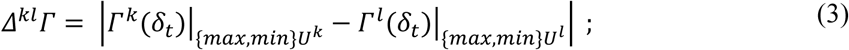

In this expression, there is an implicit difference between different landscapes of control functions originated in changes in the environment (see table 1).

Thus, these kind of systems own “ears and eyes” able to sense dynamical changes in the environment, and impose decidability on one process, since the change of one to other relative environment stops a computation, while the inherent increase of complexity is reduced. In contrast, conventional dynamical systems are centered in the computation of a single trajectory.

This also implies the existence of a family of structures, such that 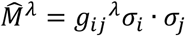 If two trajectories exist, then there is a probability to establish a distance between different structures, such that for two environments *k* and *l*, with 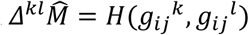 the hamming distance between these both structures.

If computational structures are deformed to explore new trajectories and make computational systems decidable, then taking elements of elasticity theory (see (Rathgeber, 2002) (Landau, 2004)), we conjecture that we can measure the deformation of these structures/mechanisms in the network. Like the elasticity modulus in solid mechanics, the elasticity modulus of mechanisms (EMM) is measured as the distance between two network structures regarding two environments 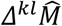 respect the mean distance between two trajectories *Δ^kl^Γ* (equation 3), and is defined as^2^

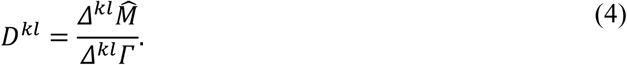

In a family of environments, and similar to the tensor notation of the elasticity modulus in solid mechanics, the expression (4) becomes a tensor-like structure:

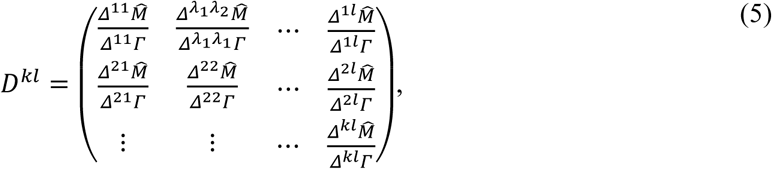

If *D^kl^* = 0 there are no changes in the structure, implying that either the system remains as non-computable, and preserves oee, or the system is complete and simple trajectories can be computed. This last case implies that the system can be mathematically described (for instance using differential equations)^3^. On the other hand, if *D^kl^* ≫ 0 there are changes in the structure 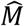 to make the system decidable. Thus, if an organism cannot compute its environment, or the trajectories are undecidable, but try to modify internal structures to meet decidable structures, then the system has been distorted.

In the next section, we present a simple example with a system exploring two different environments to illustrate the use of an elasticity modulus of mechanisms.

## Example: a modified predator-prey system with chemotactic response

Due to the fact that intrinsic networks are not consistent, there are several possible responses that can be assigned to different network architectures. To illustrate this, we employ an example for chemotaxis where is difficult *to decide* between two candidate networks associated to two different responses to stimuli (Chang and Levchenko, 2013). We argue that the system is non-computable in the sense that the system cannot decide alone which trajectory should be computed.

Here, “perfect adaptation of a signaling response turns out to be quite restrictive in terms of the number of possible ways it can be achieved. A recent analysis suggested that the ‘architecture’ or topology, of the underlying signaling networks would be expected to fall into two classes: those containing a negative (integral) feedback (NFB) and those that contain two parallel initially diverging and ultimately converging pathways, affecting the output in opposite ways. The latter network type has been termed an ‘incoherent feed-forward’ loop (iFFL)” (Alon, 2006). The, “adaptation to temporally changing inputs can be a key in this response”.

The input *z(t),* intracellular concentrations for activation and inhibition loops *(x(t)* and *y(t))* and the response *r(t)* of the organisms as well as the development of the population of consumers *C(t)* and predators *P(t),* assuming that the population behaves according to a Lotka-Volterra system, are presented in figures 2A and 2B (corresponding equations have been written down in appendix 1). The equations for the chemotactic response are adapted for the responses for a social amoeba, and were extracted from Wang et al. (Wang et al., 2012a). The results for each response are presented assuming stimuli that linearly increases with the time (Chang and Levchenko, 2013). We select a simple stimulus, and not oscillatory examples, to avoid artifacts that other kind of stimuli can introduce.

**Fig. 2A.**
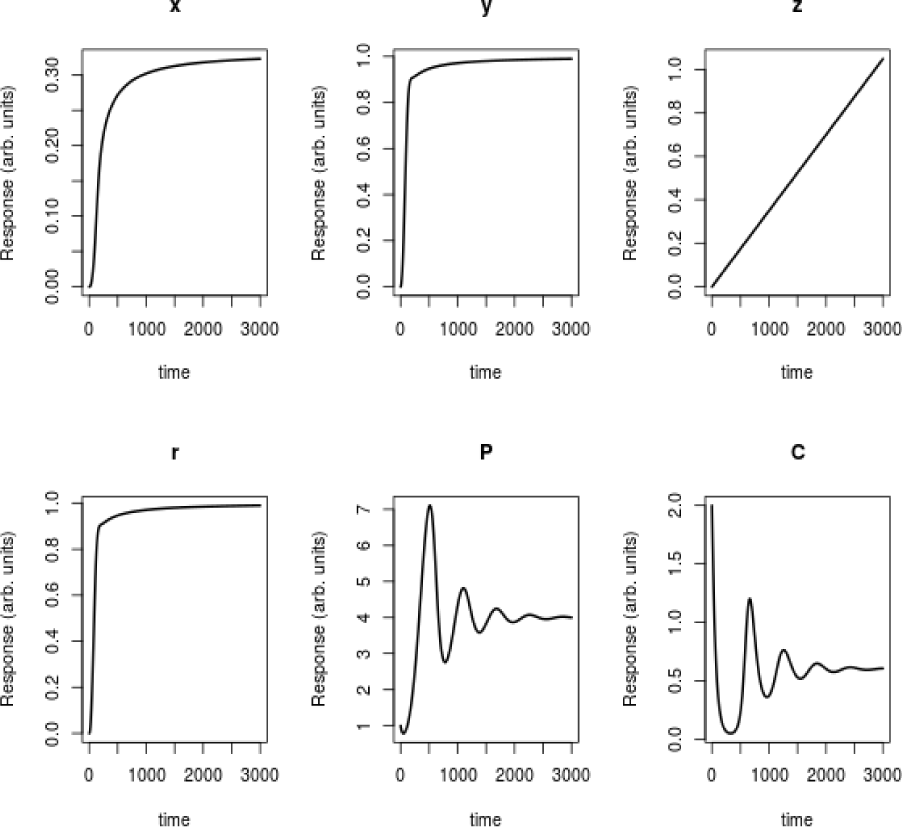
Non-adaptive response, of organisms in a population with low propagation velocity v(t). The equations describing the organism’s response, based on a NFB architecture and the population dynamics are presented in appendix 1 (equations A2)

**Fig. 2B.**
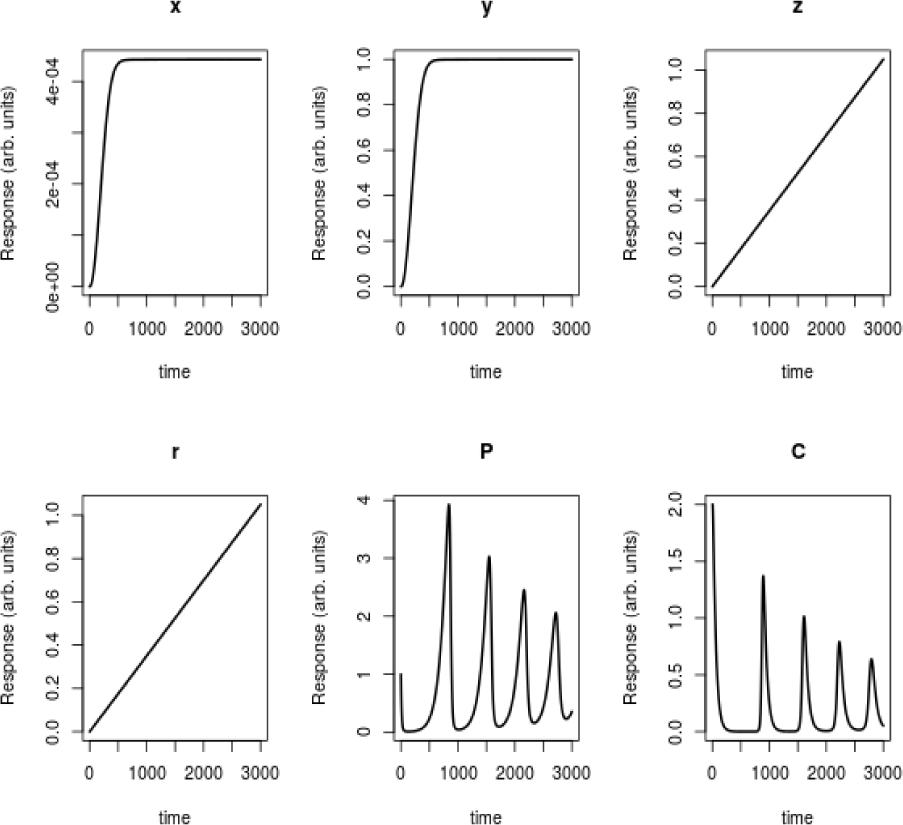
Adaptive response, in a population with high propagation velocity v(t). The equations describing the organism’s response, based on an iFFL architecture, the population dynamics are presented in appendix 1(equations A1)

The notion of distortion in this model is useful to track the completeness of the system. Additionally, to seek the reduction of the distortion helps us to define a co-evolution of internal chemo-static states in the population.

“As the rate of the change increases, the cells would tend to maximize their response, recognizing that they move in the direction leading them more precisely towards the source of the chemoattractant.” However, some organisms will require an adaptation to the external response, i.e. when they approached to the source they minimize their response, requiring the NFB but not the iFFL architecture. Here is difficult to exactly define or identify the underlying network. Furthermore, we see a system that is incomplete, i.e. two different competing models, eventually based on common molecular interactions: both can perfectly explain the behavior of the system.

We argue that both responses can belong to a cell able to adapt its response to an external source. To this end, one of both responses must be selected depending on whether the entire system (the population as organism) is incomplete or not. We do that if the cell prefers an adaptive response, only if this response lies below a threshold value. Thus, the cells as an organism “decide” (“accommodation”) which response fits better depending on the behavior of the population of consumers & predators (see figure 3). In this way, we establish a direct interaction between the single organisms and the entire population.

**Fig. 3:**
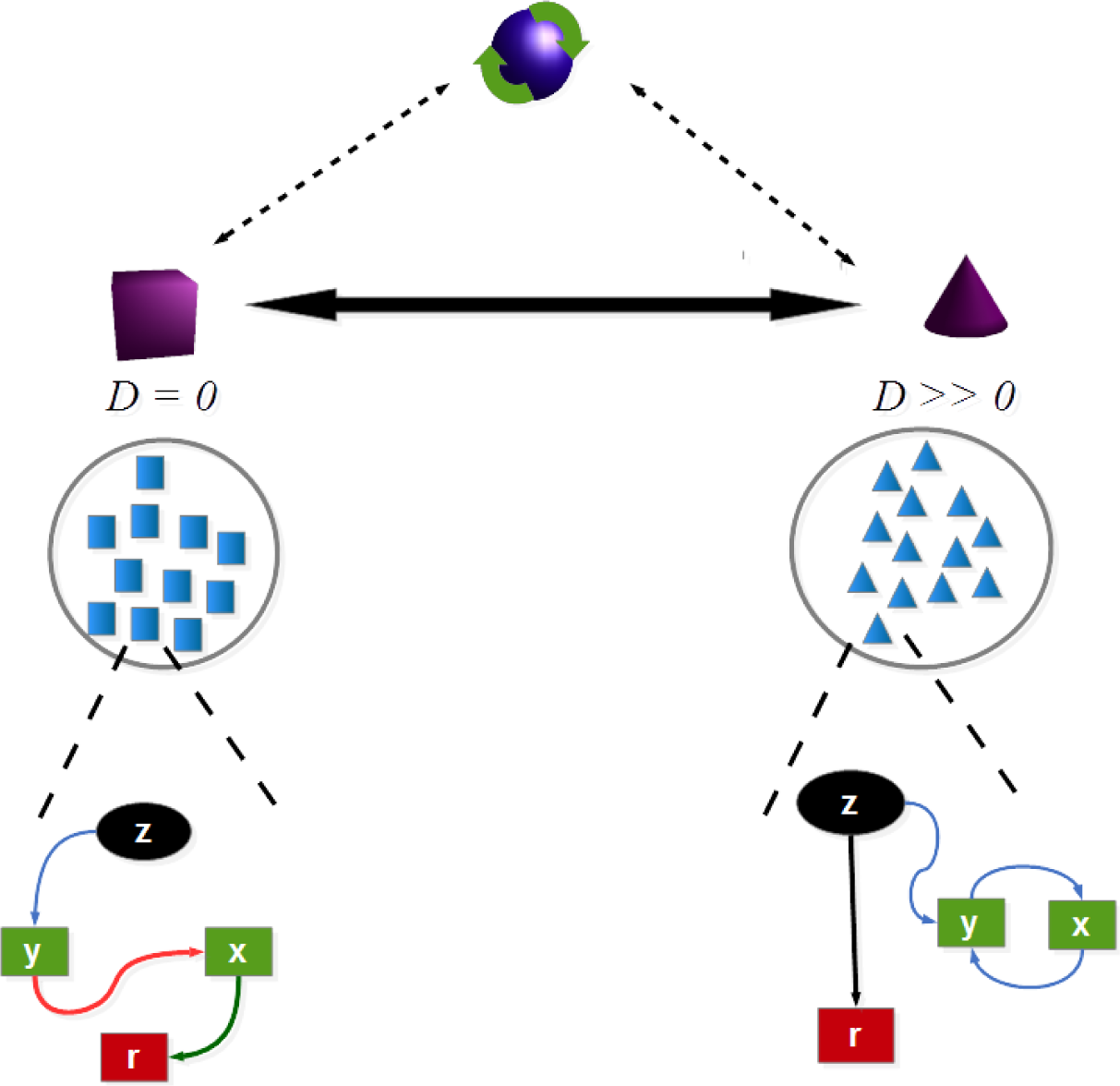
Example in chemotaxis where a population must decide between two different kind of responses: one for an *iFFL* architectureg_ij_^1^ (left - quares) and a *NFB g_ij_^2^* architecture (right - triangles), depending on the interaction of the entire population with the environment (predator’s population).

We construct a toy model using the following steps (the complete set of equations for the population dynamics are presented in equation A1 to A3 in appendix 1):

- Assume that consumers (*C*(*t*), population of predators) are close to a chemical stimulus that is frequented by a population of bacteria that is predated *C*(*t*). In our example, the stimuli growths proportional to the time.
- The bacteria have a variable response: switch from an adaptive to a non-adaptive – response. The bacteria alone cannot “decide” which is the better response; this decision is made as an organism, but accounting the entire population and its environment. The response will depend on the whole population of predators.
- If *C*(*t*), (population of predators) is relatively low, then the preferred response remains constant; otherwise, if the population of predators increases above a critical number, then the preferred response changes from a non-adaptive to an adaptive response.
- The kind of response influences the velocity towards the stimuli: a non-adaptive response is related to high propagation velocity towards the stimuli. Otherwise, an adaptive response is related to a slow propagation velocity (see experiments about changes of velocities for aggregation of bacteria in chemotaxis).

This “trick” should affect the number of predators, and should influence the cycles of the population (Arditi and Ginzburg, 1989) (figure 4).

**Fig. 4.**
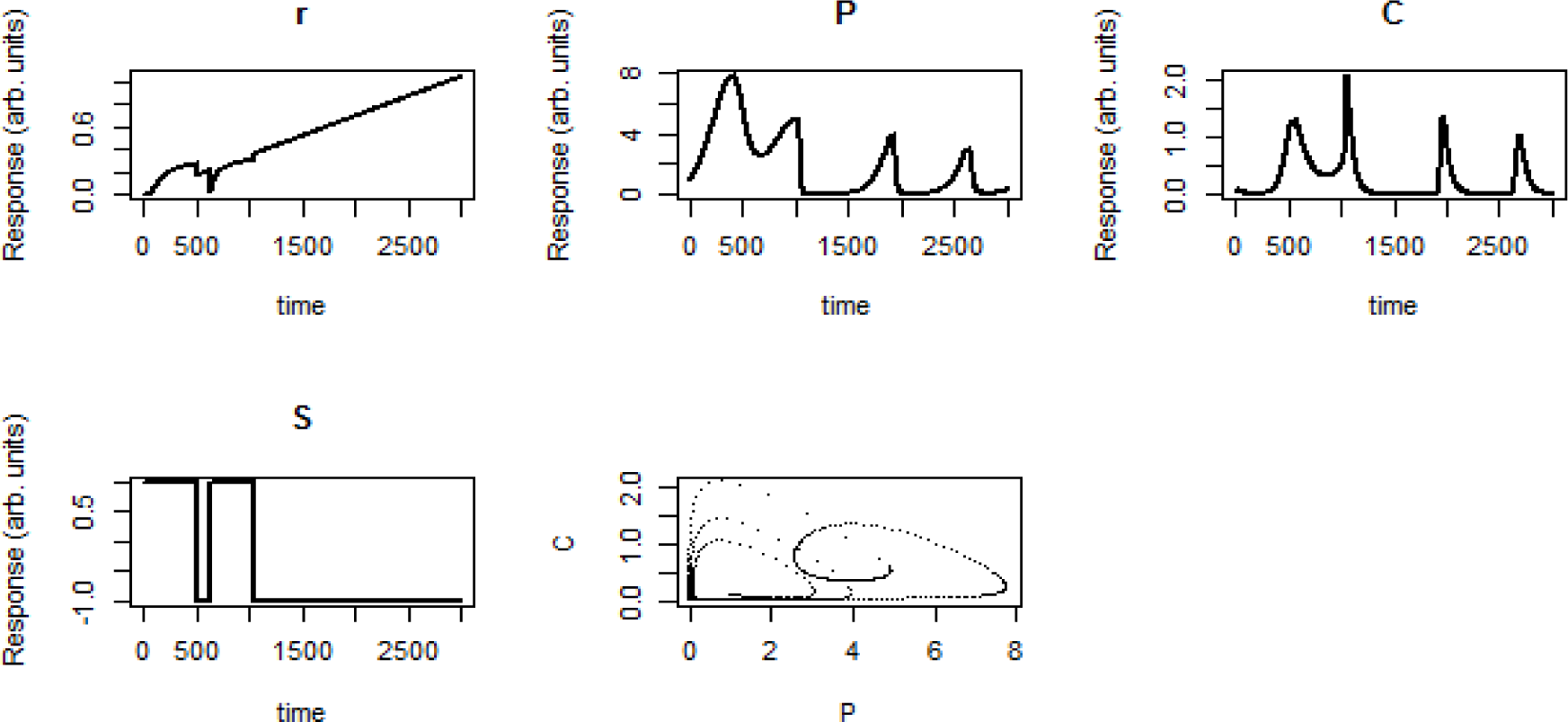
*In figure above*, *P*(*t*) *are the predated population (in our example bacteria) and C*(*t*) *are consumers. S detects which response has been selected: S = 1 is for a non-adaptive response; S =−1 adaptive response. The complete set of equations and parameters are presented in appendix 1*.

In this example, there is apparently a co-evolution. However, observe that the underlying network is incomplete since it has two potential background models that mutually compete; the apparent co-evolution is the decision to adopt one or other response depending on the pressure over the entire population of consumers and predated organisms. In this process, there is a constant distortion, until the organism meets a “decision” and selects only one response. According to equation 5, bellow the first *τ_a_* = 1000 time steps

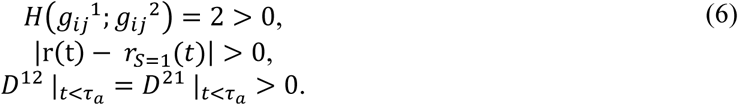

Observe that the selected parameters generate a critical behavior, in which the cells alternatively select two of the responses. The change of response depending to the predator’s population imply a distortion *D*^12^ = *D*^21^ >0 (which is known in this model). The additional oscillation of this distortion implies self-organization in the population dynamics and the adaptation of the response. Also, in this model we assume that each kind of response is complete, i.e. *D^11^* |_*t*<τ_a__ = *D^22^* |_*t*<τ_a__ = 0*^4^*.

Above a critical value of the stimuli the cell finally selects one response from the two potential responses, after 1000-time steps, implying that *D_21_* |_*t*>τ_a__ = *D_12_* |_*t*>τ_a__ =*D_11_* |_*t*>τ_a__ = *D_22_* |_*t*>τ_a__ = 0. This transition is visualized in the phase diagram in figure 4.

In this example there is uncertainty about how to choose the best response to the environment. The microstates do not simply have a passive response to the environment, but actively select a response (in our example from two possible responses), modifying their behavior at higher levels. By doing this, higher levels close the loop and generate completeness, i.e. multi-scaling is responsible for the generation of completeness.

Thus, the organism can stop or restart a computation (response or function) regarding the response of the macro-state. This also implies, that the organism can in principle maintain different redundant structures that either perform the same function or response (Tononi et al., 1999) or that can iterate different responses, a decision that depends on the multi-scaling where the organism is embedded.

## Discussion

The construction of theories requires completeness in order to make them predictive. However, we continuously experience a challenge to this completeness in various levels. Writing a text like this is an example about how difficult is to retain completeness in the transmitted message and decide when to stop. The goal is to write a code that works like a mechanism in the minds of other people; but so many ideas and concepts compete, making difficult to find a complete sentence, or decide when to stop writing. This also applies to our daily life and business, as well as biology in writing and expressing genetic codes.

In the nature we find very often incompleteness and undecidability. Several mechanisms can run to infinity, as we have shown in our vesicles example (section 2). However, the interaction with the environment, as well as the selection of functionality induces a primitive cognition that helps these processes to decide, in the frame of accommodation and assimilation (Bitbol and Luisi, 2004). Constant changes in the environment imply a change of the initial completeness, pushing living organisms to constantly try to maintain a decidability or relative completeness. This sets a limit to our ability to identify models in systems biology: while a mechanism can be valid under certain circumstances, continuous assimilation and accommodation of organisms challenge the completeness of these mechanisms. This implies a distortion of interaction mechanisms.

A measure of this distortion is therefore helpful to decide when a model is predictive or can be extrapolated. From our daily experience we know that it is almost impossible to fly by pulling on our shoe laces. Similarly, incompleteness in the nature cannot be solved from the microstates alone. Thus, “higher levels” in biology close the loop and generate completeness where a single level (the microstate) cannot, providing a framework where multi-scaling is essential for the computability of the system. In this way the distortion module quantifies the deformation of computational structures, triggered by larger scales, to explore new trajectories and make computational systems decidable.

We use this concept to model several responses in chemotaxis. If the distortion is larger than zero, then different responses associated to different interaction networks are selected depending on the population of predators. This example allows us not only to model making use of this distortion, but also to exemplify this assimilation and accommodation depending on the evolution of the populations of bacteria and predators.

However, chemotaxis is not the single field where these concepts can be applied. Cancer is also a potential candidate: “The oxygen-deprived cells (environment) suffer an excess of DNA methylation, which silences the expression of tumor-suppressing genes, thereby enabling aberrant cellular behavior and enhancing tumor growth.”^5^ (Thienpont et al., 2016). This also is related to the way how cancer is treated: while several efforts focus on the identification of biological mechanisms for the targeted treatment of the disease, the practical application has shown that this strategy often not only does not work, but is in some cases harmful to the patient. This problem is not only rooted in the complexity of the cancer mechanisms, but also on the capability of the cancer cells to evolve and develop tolerance and mutate against treatments ^6^.

Also in physics, there are potential traces of incompleteness. For instance, a toy model for spin-ice can also illustrate this incompleteness and distortion, with a connectivity of microscopic states depending on a whole energy landscape (Ochoa, 2014). It also implies that equations of life, for instance relating entropy with replication, are only valid when the energy gap of internal reservoirs respect the environment is zero (England, 2012).

These facts are against the definition of laws for complex systems. Perhaps we can identify rules; but rules are not laws, which is a fact that start to be recognized not only in biology but in general in many social and technological system that my “require removing the segregation of states and *fixed* dynamical laws characteristic of the physical sciences for the last 300 years” (Adams et al., 2017). This fact allow us to observe regularities that cannot be generalized for all the systems, in the sense of a strict generalization or at least as a *ceteris-paribus* rule (Carroll, 2016). This is true not only for biology, but also for social sciences. Mathematically this has a deep implication: whereas in physical systems it is possible to recognize laws and fundamental models that in principle work in every part of the universe, for complex systems there are rules, rather than laws, that are often subject to exceptions.

Mathematically this also implies that there is no methodology to produce good universal predictions, and that biology cannot be solely described by using differential equations (Danchin, 2009). This is only valid when *D_kl_* = 0. Once we want to predict something we are confronted to the necessity to constantly collect and update information.

This also implies that when using networks to describe mechanisms certain mathematical laws must be considered more as a rule that can be subjected to exceptions. For example, the concept of scale-free distribution of node connectivity in networks is perhaps a rule (Barabási, 2009), but not a law, which can be continuously challenged by the incompleteness of molecular networks (Khanin and Wit, 2006).

Finally, in its current form the measurement of the distortion of the mechanisms with an elasticity modulus should not be confused with the distortion-rate theory, initially defined by Claude-Shannon and which has been applied in recent works in biology by Marzen and DeDeo to study the “evolution of lossy compression and quantify the trade-off between acquisition costs and perceptual distortion, allowing to talk about the extent to which an organism can save costs of transmitting a representation of their environment by selectively discarding information” (Marzen and DeDeo, 2017).

Both approaches, the EMM and the distortion rate theory of Marzen and DeDeo, handle two different aspects: the first focus on the computability of dynamical systems; the second is essentially an optimization problem that quantifies the optimization of acquisition costs respect to perceptual distortion. However, in their roots, both information theory and computation theory are deeply interlinked (Danchin, 2009), implying that a more detailed study is required to link both concepts in a single theoretical framework.

## Concluding remarks

This work is an alternative to the conventional bottom-up or up-bottom approaches to comprehend living systems. In our opinion, there is not a hierarchy of scales, but much more an interaction across different scales, as has also been pointed by Ellis for instance as top-down causation in adaptive selection (Ellis, 2012).

Incompleteness is also the impossibility to know how an organism and its mechanisms has specific functions regarding evolutionary pressures. Certainty there are well defined functions, but organisms across scales fulfill so many tasks, that as a result is difficult to make optimal definitions. As we have shown in our toy model, it is useful to consider contrasting objective functions and try to obtain one solution by accepting that the organism decides which function will adopt depending on how consistent this objective function is with respect to its environment.

Then, can biology or complex social and technological systems be explained using fundamental theories or even physics? Our answer is: only sometimes. Occasionally we can define laws that envision the possibility to describe these systems by means of fixed dynamical laws like in physics (Goldenfeld and Woese, 2011), but in general it is much more cautious to speak about rules. We also recognize that in these kind of systems rules are non-invariant (an opinion currently shared by other authors (Adams et al., 2017)). This position is not a pessimistic statement, but is instead a stimulating chance, that is coupled to our current technological ability to increase measurements and use stream data and artificial intelligence to better understand and deep in the knowledge of evolving systems, particularly in systems biology.

## Acknowledgements

I want to acknowledge the constructive input from two anonymous referees who helped me to refine and set this work on a solid basis. Without this collaborative work with the reviewers it hasn’t be possible to give the final form of this manuscript. I am also grateful with Elena Ramírez for decisive discussions that provided the fundamental elements exposed in this manuscript.

## Footnotes

1 This adaptation explains why the rolling stones have survived so many time despite drug and general health abuse.

2 This relation is valid since we relate two distances. No additional mathematical structures are here involved.

3 Both conditions are dangerous, since dynamical systems could be used to describe systems that are oee. This can be wrong if the existence of the gap between trajectories describing the environment is not recognized.

4 Otherwise we must assume for instance more than one adaptive response associated to different models.

5 http://www.genengnews.com/gen-news-highlights/cancers-grow-by-throwing-epigenetic-smother-parties/81253107/

6 http://www.nature.com/nature/journal/v537/n7619_supp/full/537S63a.html

